# Leveraging Machine Learning-Guided Molecular Simulations Coupled with Experimental Data to Decipher Membrane Binding Mechanisms of Aminosterols

**DOI:** 10.1101/2024.01.31.578042

**Authors:** Stefano Muscat, Silvia Errico, Andrea Danani, Fabrizio Chiti, Gianvito Grasso

## Abstract

Understanding the molecular mechanisms of the interactions between specific compounds and cellular membranes is essential for numerous biotechnological applications, including targeted drug delivery, elucidation of drug mechanism of action, pathogen identification, and novel antibiotic development. However, the estimation of the free energy landscape associated with solute binding to realistic biological systems is still a challenging task. In this work, we leverage the Time-lagged Independent Component Analysis (TICA) in combination with neural networks (NN) through the Deep-TICA approach for determining the free energy associated with the membrane insertion processes of two natural aminosterol compounds, trodusquemine (TRO) and squalamine (SQ). These compounds are particularly noteworthy because they interact with the outer layer of neuron membranes protecting them from the toxic action of misfolded proteins involved in neurodegenerative disorders, both in their monomeric and oligomeric forms. We demonstrate how this strategy could be used to generate an effective collective variable for describing solute absorption in the membrane and for estimating free energy landscape of translocation via On-the-fly probability enhanced sampling (OPES) method. In this context, the computational protocol allowed an exhaustive characterization of the aminosterols entry pathway into a neuron-like lipid bilayer. Furthermore, it provided accurate prediction of membrane binding affinities, in close agreement with the experimental binding data obtained by using fluorescently-labelled aminosterols and large unilamellar vesicles (LUVs). The findings contribute significantly to our comprehension of aminosterol entry pathways and aminosterol-lipid membrane interactions. Finally, the deployed computational methods in this study further demonstrate considerable potential for investigating membrane binding processes.

## Introduction

Cell membranes play an essential role in all cells, separating the intracellular environment from the extracellular milieu as well as a vast array of internal compartments. The presence of a hydrophobic core and the compact arrangement of lipid components create semi-permeable barriers ensuring the cell homeostasis^1^. The translocation of molecules across cellular membranes is accomplished via both active and passive permeation strategies. Active transport involves energy consumption to move molecules against their concentration gradient, generally facilitated by transport proteins. In contrast, passive transport, which includes the protein-indipendet simple diffusion and protein-mediated facilitated diffusion, operates independently of energy, facilitating the movement of molecules along their concentration gradient. The above-mentioned mechanisms are crucial for various physiological processes. For example, they mediate the exchange of O_2_ and CO_2_ across the membrane of erythrocytes, supporting cell signalling by allowing second messengers like H_2_S to reach their respective targets and pathogen detection^1–4^. In the pharmaceutical domain, key stages as gastrointestinal absorption and portal venous system crossing, which primarily rely on trans-cellular diffusion, are of paramount importance^5,6^. The identification of new compounds with the ability to penetrate lipid membranes is fundamental toward the design of novel inhibitors of intracellular target. Despite the central role of strong molecular binding in securing drug efficacy, its potential can be thoroughly undermined when poor membrane permeability hampers the bioavailability in live organisms^7^. Moreover, recent findings on G protein-coupled receptors (GPCRs), the primary targets for approximately 33% of all small-molecule drugs currently available^8,9^, propose that amphiphilic and lipophilic molecules may interact with these receptors by initially partitioning into the membrane. These molecules reach the binding site via lateral diffusion along the lipid bilayer^10^. In this context, the lipid bilayer can enhance ligand-receptor binding efficiency, even at low concentrations, by confining the drug within a particular bilayer region. However, while membrane interactions can enhance binding kinetics, excessive membrane accumulation can cause toxicity due to off-target interactions, making the simple increase of ligand lipophilicity undesirable. Comprehensive understanding of factors such as bilayer distribution, preferred location, orientation, and conformation of drug molecules within the bilayer is crucial for understanding their target binding kinetics, onset and duration of action, and disposition^10–12^. Recent breakthroughs in experimental methods have yielded a wealth of data on membrane interactions of diverse molecules^13,14^. However, elucidating the detailed mechanisms of drug insertion into membranes, including their orientation and location, remains an intricate challenge. Molecular dynamics (MD) simulations can supplement this shortcoming, offering quantitative, atomic-level insights into chemical interactions with lipid bilayers, thereby enriching the experimental data^15–18^. However, classical MD simulations often fail to accurately capture rare events such as passive membrane permeation. The free-energy barrier that a compound needs to surmount to diffuse into cell membranes, further exacerbates this problem by restricting the compound to a narrow range of configurations around a starting state. Therefore, several enhanced sampling methods have been developed to sample inaccessible configurational spaces. Among the most widely used advanced sampling methods is umbrella sampling, where a solute is restrained with respect to specified locations along a lipid bilayer^19,20^. Furthermore, simulated tempering-enhanced umbrella sampling^21^, replica exchange MD (REMD)^22^, metadynamics (MetaD)^23^ and transition tempered MetaD^24^ and other variants have been applied in solute-membrane absorption studies^25,26^.

A challenging aspect of these advanced methods is the proper selection of one or more reaction coordinates or collective variables (CVs). Identifying the optimal CV stands as a considerable challenge given that an intuitive articulation of the process might overlook critical orthogonal degrees of freedom inherent to it. Enhanced sampling techniques face challenges in exploring intricate system dynamics with few physical collective variables. Effectively sampling a complex system may require up to hundreds of CVs^27^. The design of CVs, which seeks to uncover a low-dimensional representation from high-dimensional configurations, aligns with the goals of dimensionality reduction algorithms. These techniques rely on the assumption that data points exist near a low-dimensional manifold, even when situated in a high-dimensional space. For this reason, a variety of data-driven methodologies and signal analysis methods have been presented as potential avenues for CVs creation^28^. One such method, Time-Lagged Independent Component Analysis (TICA), is a linear transformation technique that identifies coordinates exhibiting maximal autocorrelation at the specified lag time. The driving principle behind TICA is rooted in the idea that optimal CVs ought to represent the slow modes of a molecular system, as these modes exhibit correlation functions that decay slowly over time. The integration of TICA methodology with MetaD has been investigated in literature, showing diffusive phenomena within the domain of the CV^29–31^. Recently, the effectiveness of linear methods in identifying CVs has been significantly enhanced by incorporating Neural Networks (NNs), which capitalize on their capacity to approximate non-linear functions of multiple variables^32^. This integration has resulted in the development of highly efficient CVs, as demonstrated by innovative approaches like the reweighted autoencoded variational Bayes for enhanced sampling (RAVE)^33^ and Deep-TICA^32^. These methods harness the power of deep learning to better capture the complex relationships and dynamics within molecular systems, leading to improved sampling performance and a more accurate representation of the underlying free energy landscapes^34^.

Here, we present a novel application of an established computational framework for calculating the free-energy surface in solute-membrane insertion processes, employing a coarse-grained (CG) modeling approach. The framework is based on the synergistic combination of enhanced sampling methodologies and Deep-TICA^32^. This protocol leverages the On-the-fly probability enhanced sampling (OPES) method, a recent advancement in the MetaD methodology, as its foundational element^35,36^. Moreover, we validated the effectiveness of our protocol by predicting the membrane binding affinity of two natural aminosterols, namely trodusquemine (TRO) and squalamine (SQ)^37,38^. These aminosterols are known to interact with the outer membrane of neurons, providing protection from the toxic effects of amyloidogenic proteins, in their monomeric or oligomeric forms, which are linked to neurodegenerative diseases like Alzheimer’s (AD) and Parkinson’s (PD)^15,17,37,39–42^. In this context, the computational protocol allowed a comprehensive characterization of the entry pathway of aminosterols into the lipid bilayer. The predicted aminosterol-membrane binding affinity have been validated using experimental fluorescence emission techniques involving BODIPY TMR-labeled aminosterols in presence of large unilamellar vesicles (LUVs) with the same lipid composition used in MD simulations. This integrated approach underscores the robustness of our methodology and paves the way for deeper investigations in the solute-membrane insertion process.

## Materials and Methods

### Simulations set up

In this study, we focused on the interactions between a lipid bilayer and two distinct molecules: TRO and SQ (Figure 1). More specifically, a membrane composition of 200 lipids was symmetrically modelled using the python tool insane^43^, composed by 59% of 1,2-dioleoyl-sn-glycero-3-phosphocoline (DOPC), 30 % of sphingomyelin (SM), 10 % of cholesterol (CHOL), and 1% of monosialotetrahexosylganglioside 1 (GM1). We employed the polarizable water model from the Martini 2.2p force field for solvating the lipid bilayer and incorporated 150 mM NaCl to neutralize the net system charge. Finally, each complex consisted of the lipid bilayer, with the aminosterol molecule in the water environment, for a total of about 20,000 interacting particles. The MD simulations were performed with GROMACS 2021 software package patched with PLUMED 2.9^44^ and the Pytorch library 1.4^45^. Each molecular complex was first energetically minimized. To equilibrate each system, 1 ns in NVT ensemble at 310 K and 5 ns in NPT ensemble at 1 bar and 310 K simulations were sequentially performed starting from each initial minimized structure. The temperatures and pressures of all systems were controlled using v-rescale thermostat^46^ and Parrinello−Rahman barostat^47^, respectively. The electrostatic interactions were calculated using the particle mesh Ewald method^48^ with a real space cut-off of 11 Å. The cut-off value for van der Waals interactions was set at 11 Å.

**Figure 1:**
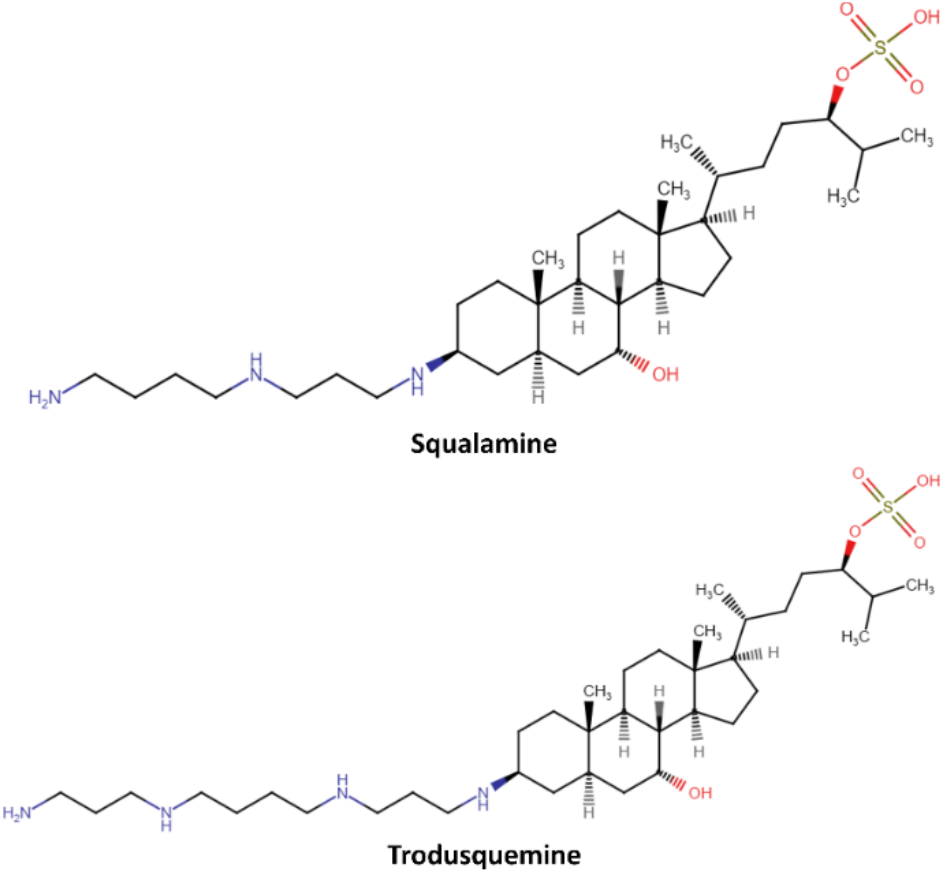
SQ and TRO chemical structures. Both aminosterols are formed by a sulphate group (right) and a sterol (centre). The structural distinction between SQ and TRO lies in the polyamine moiety connected to carbon-3 (C-3) of the molecule (left). In SQ, the polyamine attached to C-3 is spermidine, while in TRO, it is spermine.

### Martini Force Field

The Martini 2.2p force field^49^ was used to represents the molecular components. The force field concept is based on a four-to-one mapping, in which a single interaction center named ‘bead’, represents four heavy atoms and their associated hydrogens. Each bead has a number of subtypes, allowing for an optimal balance between computational efficiency and chemical representation. The three-bead polarizable water bead was employed to correctly reproduce the orientational polarizability of real water^50^ and the ions were represented by a single CG bead. The aminosterols mapping was obtained from an automated approach that preserve the symmetry of the molecules, and correctly reproduce the experimental octanol-water and water-membrane partitioning data^51^. Finally, the bond terms of TRO and SQ molecules were parametrized employing swarm-cg tool^52^ (see Supporting Information S1 section, Figure S1-S6).

### OPES Expanded and OPES MetaD

To enhance the sampling the OPES method was employed^35^. OPES belongs to adaptive bias methods and is a development of MetaD^53,54^ in which Gaussian kernels are utilized to rebuild the marginal probability distribution along the CVs as opposed to directly creating the bias potential. The equilibrium probability distribution is estimated on the fly and the bias is selected to drive the system toward a target distribution. By selecting the proper target distribution, one may generate random samples from a wide range of ensembles, including the well-tempered^53^ or a generalized ensemble^36^. In this work, two different distributions are targeted. In a first set of simulations, the multithermal distribution is sampled to allow rapid exploration of the thermodynamic states accessible to the system through the OPES expanded method^36^. In detail, a temperature range between 270K and 500K was explored. For every individual molecular system under study, eight replicas of 1 μs that shared the same bias potential, were conducted, accumulating a total simulation time of 8 μs. Next, the well-tempered distribution^36^ is sampled using the deep-TICA-1 vector as CV employing the OPES MetaD method. An adaptive kernel width was employed, setting the minimum value that can be achieved to 0.028. The maximum energy barrier that can be overcome was established at 60 kJ mol^-1^, and the deposition rate was set to occur every 500 simulation steps. For each molecular system, 8 replicas of 400 ns each were simulated for a total of 3.2 μs of simulation time. In addition, the replicas shared the same bias potential in order to harvest more transitions between the ligand’s states in and out the lipid bilayer.

### Deep-TICA

The recent Deep-TICA method^32^ was employed to design a comprehensive CV for membrane insertion process. Deep-TICA is based on the TICA that aimed to solve the VAC through a linear solution. For a comprehensive theorical background of Deep-TICA, readers are referred to recent studies by Bonati et al^32^. The Deep-TICA CV has been trained on the converged OPES expanded multithermal simulation of each system using the PyTorch library. The first 200 ns were discarded and a set of 180 (TRO system) and 168 (SQ system) molecular descriptors were evaluated during the remaining simulation time. A feed-forward NN was employed, composed of an input layer with 180 and 168 nodes for TRO and SQ, respectively. Furthermore, two hidden layers containing 128 and 64 nodes were added. The hyperbolic tangent was used as activation function. The dataset was split into training/validation sets. To optimize the neural network parameters, we used the ADAM optimizer with a learning rate of 1e-3. The lag time was set equal to 0.01 and in order to prevent overfitting the early stopping with a patience of 10 epochs was employed. Additionally, we scaled the inputs to have a zero mean and a variance of one. The Deep-TICA CVs were also adjusted, ensuring their value range fell between -1 and 1. Finally, the trained model was exported as serialized model to be exploited during the OPES MetaD simulations.

### Preparation of Large Unilamellar Vesicles (LUVs)

Liposomes were produced with the same lipid mixture of the computational model: 59% (mol) DOPC (Avanti Polar Lipids), 30% (mol) SM (Sigma-Aldrich), 10% (mol) CHOL (Sigma-Aldrich), and 1% (mol) GM1 (Avanti Polar Lipids). The lipids were dissolved in chloroform/methanol (2:1), and the organic solvent was subsequently removed by evaporation in vacuo (Univapo 150H, UniEquip) for 180 min. The mixture was hydrated at a total lipid concentration of 2.0 mg/ml with distilled water to form multilamellar vesicles (MLVs), left to swell for 1 h at 60 °C, and then extruded 17 times through a polycarbonate membrane with 100 nm pores using a miniextruder (Avanti Polar Lipids) at the same temperature, to form large unilamellar vesicles (LUVs). After cooling to room temperature, LUVs were stored at 4 °C for a maximum of 1 week.

### Labeling of TRO and SQ with BODIPY TMR

SQ and TRO were synthetized by coupling spermidine and spermine, respectively, to the (5α,7α,24R)-3-keto-7-hydroxycholestan-24-ol sulfate steroid intermediate as previously described^55–57^, and stored as powders until use. For the labeling procedure, the two aminosterols were dissolved in distilled water to obtain a 100 mM stock solution and stored at 4 °C. BODIPY TMR-X NHS Ester (BODIPY TMR, ThermoFisher Scientific) was dissolved in DMSO to obtain a 15 mM stock solution, and stored at −20 °C. For labeling, 5 mM AMs, 0.5 mM dye, 0.1 M sodium bicarbonate buffer, pH 8.3 were incubated in a final volume of 15 μL at 25 °C for 3 h under mild orbital shaking. During labeling procedure, TRO and SQ precipitate. Therefore, after the incubation, the solution was centrifuged at 18,000 g for 15 min; the pellet was dried with a nitrogen flow and resuspended in 20 μL DMSO to maintain the initial concentrations. The labeled:total aminosterol was 1:10, no unreacted dye was detected using mass spectrometry, following a previously described procedure^17^. As a negative control, L-Arg was labeled with BODIPY TMR under the same conditions used for aminosterol labeling and no precipitate was observed.

### Binding assay of BODIPY TMR-labeled TRO and SQ and LUVs

BODIPY TMR-labeled TRO, SQ and L-Arg (negative control) were diluted with distilled water to a final concentration of 10 μM and incubated with increasing concentrations of unlabeled LUVs composed as described above (from 0.0 to 1.0 mg/ mL) for 15 min at 25 °C in the dark. Fluorescence emissions of BODIPY TMR-labelled species were then acquired at 572 nm after exciting the samples at 535 nm, using a 3 × 3 mm black walls quartz cell at 25 °C on an Agilent Cary Eclipse spectrofluorometer (Agilent Technologies) equipped with a thermostatted cell holder attached to an Agilent PCB 1500 water Peltier system. The resulting emission values were normalized to the value obtained in the absence of LUVs (taken as 100%) after the subtraction of unlabelled LUVs contributions and then plotted versus LUV concentration. Data points were then fitted with:

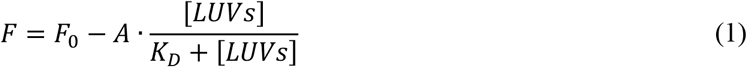

where *F* is the fluorescence intensity at a given LUV concentration, *F*_*0*_ is fluorescence intensity in the absence of LUVs, *A* is the difference between the fluorescence emission of unbound and bound aminosterols, and *K*_*D*_ is the dissociation constant of the LUV-aminosterol complex.

## Results and Discussion

### Design of collective variables for aminosterols insertion into phospholipid bilayer

The membrane insertion of two natural aminosterols, TRO and SQ was investigated in this study by leveraging enhanced sampling techniques in combination with machine learning algorithms, applied to address the challenge of estimating the free energy associated with the membrane insertion process of small molecules. These natural aminosterols, initially discovered and identified in the digestive tracts of dogfish sharks (*Squalus acanthias*)^37,40,58^, represent a promising prospect in the search for effective treatments against AD and PD^40,42,59,60^. From a structural viewpoint, TRO and SQ are very similar (Figure 1). Both are cationic amphipathic aminosterols composed of a sulphate moiety, a central sterol group, and an alkyl polyamine group on the other side of the sterol: TRO consist of a spermine moiety, while SQ has a spermidine moiety. Recent evidence showed the ability of these two molecules to influence self-assembly kinetics of the amyloidogenic proteins responsible of AD and PD, displacing them from the cell membrane^39,42^. In addition, both aminosterols demonstrated to interact with the neuronal cell membrane, thereby offering protection against the toxicity induced by oligomers responsible of AD and PD^15,17,39–42,61,62^. In particular, they were shown to change the lipid distribution within the membrane bilayer, to increase the mechanical resistance force, and to reduce the natural membrane negative charge^17,63^. Examining the molecular interactions between the aminosterol molecules and the cell membrane contributes to a deeper understanding of the aminosterol molecular mechanism of action. This knowledge is critical for the development of effective pharmacological interventions and the optimization of treatment outcomes. In the drug discovery pipeline, accurately estimating ligand-binding affinity is crucial as it facilitates various steps, including structure-based drug design and lead optimization.

The binding affinity of the aminosterol molecules can be more precisely estimated through the application of enhanced sampling methods that leverage CVs to simplify the complexity of the molecular system and allow a more detailed exploration of molecular interactions and transitions. Two are the primary approaches to constructing an effective CV. The first approach requires a deep understanding of the molecular system, enabling an expert to identify a few representative variables for the system. However, the proper sampling of such complex systems that describes the interplay between a number of molecular players may require up to hundreds of CVs^27^. On the other hand, we could collect a number of transitions between the two metastable states and compute specific molecular descriptors capable of distinguishing these states. Within this framework, the implementation of machine learning could enable us to discover the non-linear combination of CVs that aptly depict the molecular transitions, drawing from those initially identified through enhanced sampling techniques^32,33,45^. In the present study, we utilized OPES expanded targeting a multithermal distribution to observe a number of transitions between aminosterol in the aqueous environment and aminosterol absorbed in the membrane. The OPES expanded multithermal simulation enables to observe a certain number of transitions involving aminosterols absorption into the membrane (Figure S7A, C, Supporting Information), similar to the rough transitions observed in previous studies that applied a well-tempered MetaD protocol^64^. By manipulating the system temperature, multithermal simulations inherently accelerated the system dynamics, reducing the energy barriers separating the metastable states. However, in complex molecular systems, this approach might not be sufficient to observe numerous transitions and accurately estimate the free energy profile. Therefore, it becomes essential to construct a CV that aptly describes the phenomenon of aminosterol absorption in the membrane.

In solute translocation simulations across lipid bilayers, the commonly employed CV is the relative center of mass displacement between the solute and the membrane along the membrane normal usually identified with the Z coordinate^7,65,66^. However, this CV exhibits a limitation in its inability to expedite the convergence of orthogonal degrees of freedom, which include parameters such as the solute orientation, interaction patterns between solute and lipid, and lipid-lipid arrangements^21,67^. An approach to design an effective CV involves utilizing TICA, which focused on identifying the most slowly decorrelating modes, through the variational principle^68,69^. The variational principle leads to a generalized eigenvalue equation when the modes are expressed as a linear combination of descriptors^70^. In this context, Deep-TICA^32^ is a combination of TICA with hidden layers of a feed-forward NN enables its application beyond just linear combinations of descriptors, significantly improving the variational flexibility of the solution and enhancing its overall quality^71^.

Utilizing data from the OPES expanded multithermal simulations, we aimed to construct a CV capable of distinguishing between the aminosterol in a water environment and the aminosterol inserted into the lipid bilayer. To achieve this goal, each molecular system was evaluated using a series of descriptors. These descriptors were computed based on the minimum distance between each ‘bead’ in the aminosterol molecule under investigation and the group of ‘beads’ equal to each other in each phospholipid type considering DOPC and SM. For the two aminosterols studied, we evaluated a total of 180 descriptors for TRO and 168 descriptors for SQ. The descriptors were used as input for the Deep-TICA method, which combined the benefits of TICA with the flexibility and adaptability of a neural network (Figure 2).

**Figure 2:**
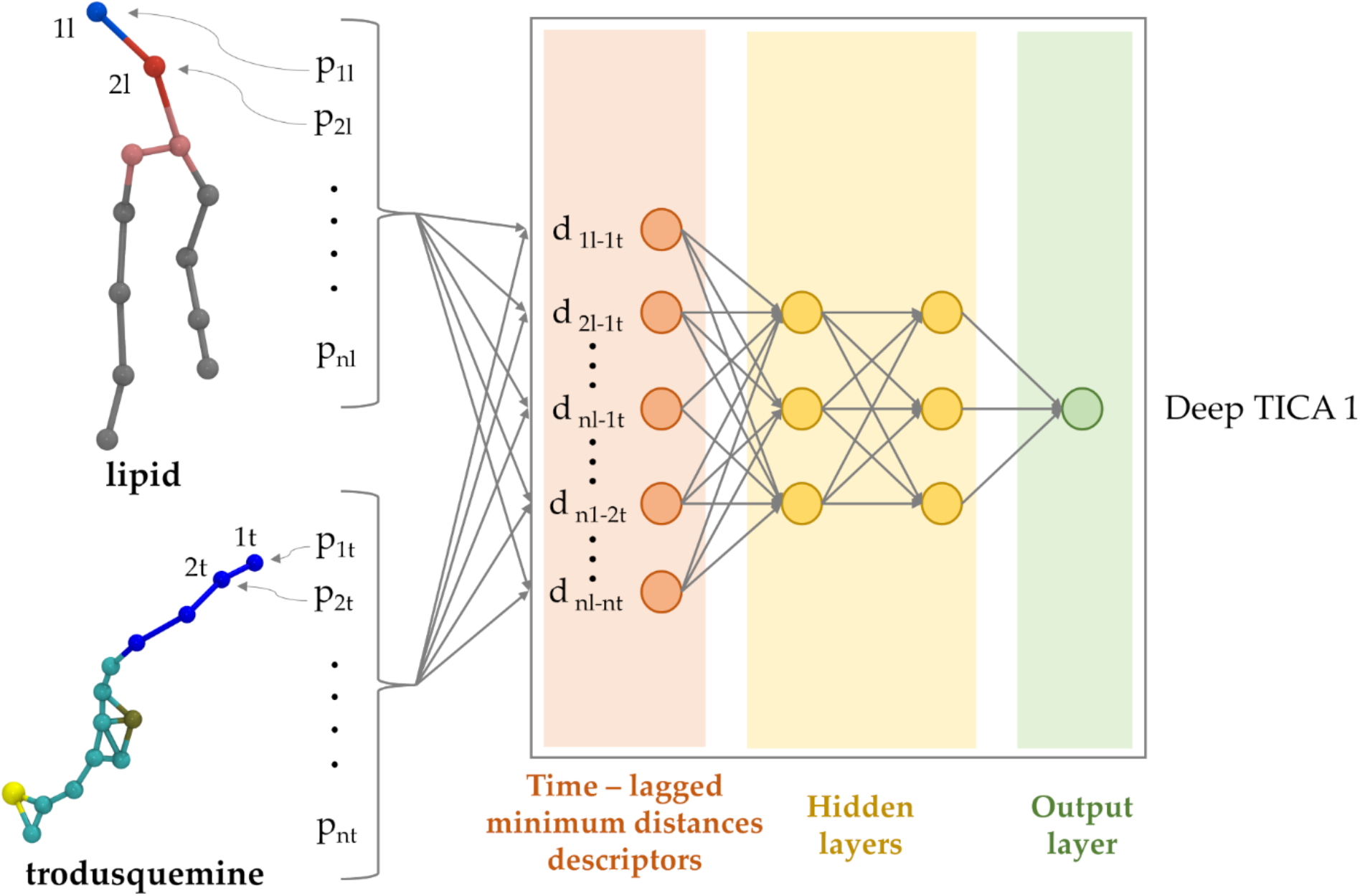
Constructing the Deep-TICA CV. The minimum distance between each ‘bead’ of the aminosterol and a group containing equal ‘beads’ from the lipids is calculated for each molecular system. These descriptors serve as the input layer of a neural network with two hidden layers, containing 128 and 64 neurons respectively. The output layer represents the Deep-TICA CV.

In each molecular system, the loss function was optimized with respect to four eigenvalues (Figure S8, S9, Supporting Information). The obtained Deep-TICA CV accurately framed the differences between the aminosterol present in the aqueous milieu and the aminosterol inserted into the lipid bilayer. Figure 3 presents a two-dimensional scatter plot illustrating the relationship between the Z component of the aminosterol dipole moment and the Z distance to the membrane, measured from the aminosterol center of mass to the membrane center. This relationship is depicted as a function of the Deep-TICA 1 CV value, which is assessed during the OPES expanded multithermal simulations. As a result, the developed CV successfully described the two metastable states of the molecular system under investigation. Specifically, the two metastable states of interest were located at the extremes of Deep-TICA 1 CV. When the CV is equal to +1, the aminosterol molecule was located in the aqueous environment. In contrast, when the CV was equal to -1, absorption into the lipid bilayer occurs.

**Figure 3:**
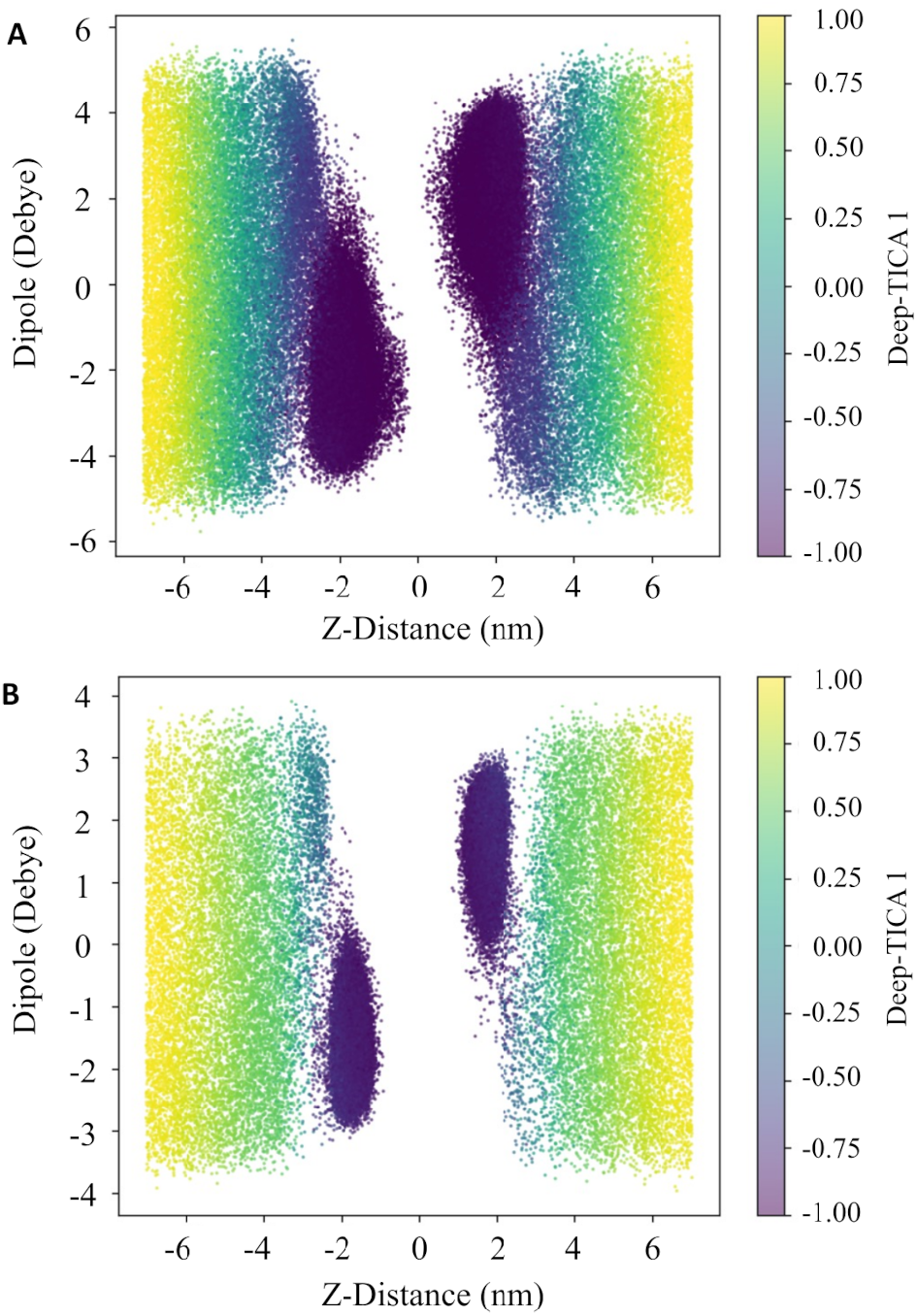
Validation of the Deep-TICA CV. Scatter plot representation of the ability of the developed Deep-TICA CV to differentiate the distinct metastable states of aminosterols (TRO panel A, SQ panel B) when interacting with the cell membrane. The plot showcases the Z component of the aminosterol dipole moment and the perpendicular distance between the aminosterol center of mass and the membrane center, colored by the Deep-TICA 1 value.

### TRO and SQ thermodynamic properties prediction

Unbiased MD simulations struggle to build an accurate free energy surface (FES) due to the limited accessible time-scales and the rarity of membrane penetration events. For a compound to enter cells through diffusion, it must overcome a significant free-energy barrier present in the cell membrane. The presence of this barrier acts as a roadblock, limiting the MD simulation ability to thoroughly sample the large configuration space. This limitation highlights the need for enhanced sampling techniques and constructing suitable CVs to better understand the molecular behavior and overcome these challenges in simulating complex systems.

In this context, we employed OPES MetaD to enhance the absorption of TRO and SQ to the membrane, leveraging the developed Deep-TICA 1 variable as CV. In Figure 4, we presented the FES in a physically interpretable space characterized as a function of the normal distance from the aminosterol and the membrane center and the normal component to the dipole moment of the aminosterol. The analysis revealed two main states, separated by a free energy barrier. These states corresponded to the basins found in water (state A) and lipid (state B) environments. In addition, the free energy difference between the A and B states was greater for the SQ molecule, indicating a greater affinity for the neuron-like membrane.

**Figure 4:**
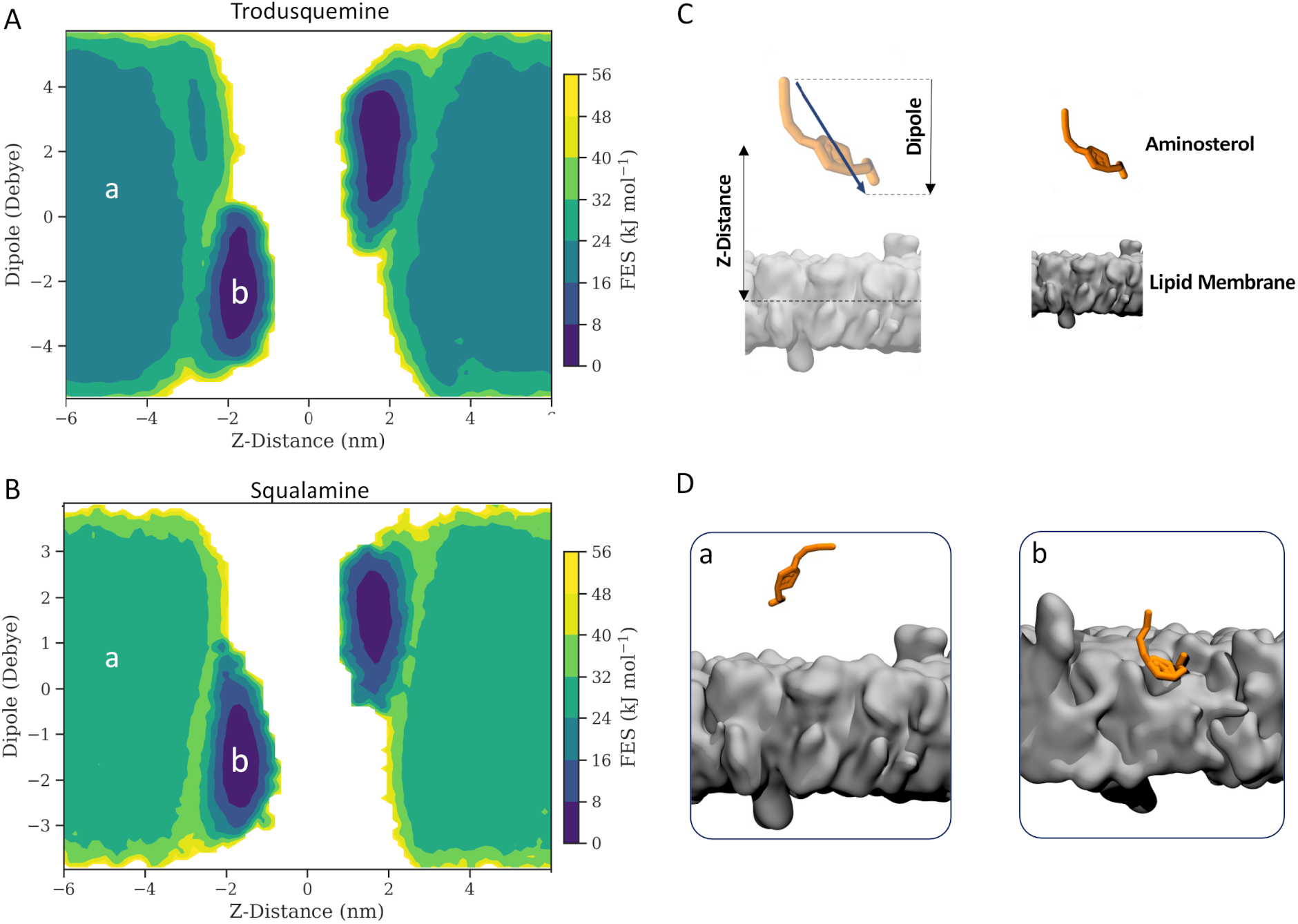
FES of aminosterols absorption. The FES of A) TRO and B) SQ are illustrated as a function of the Z-distance from the membrane center and the Z-dipole moment. C) Schematic representation of the z-distance and dipole moment CVs for both aminosterols. D) FES minima representing the metastable states, are depicted: a) aminosterols in the water environment, and b) aminosterols absorbed to the lipid bilayer.

The orientation of the molecule in the membrane played a crucial role in describing the energy minimum. This was achieved by considering the Z-component of the dipole moment of aminosterols, which encompasses information about the molecule orientation and charge. Analysis of the orientation of both aminosterols resulted in determining an angle of about 55° for the major axis of the molecule with respect to the normal to the bilayer plane, in agreement with previous unbiased atomistic MD simulations^17^. However, the dipole moment was not an appropriate variable for understanding aminosterol affinity for the lipid membrane, as identical dipole moment values can correspond to significantly disparate FES values. Instead, we considered the distance along the membrane component as a representative variable for the free energy difference (ΔG) between the state A and B. Figure 5A showed the FES along Z-Distance, where TRO exhibited a ΔG value of -16.77 ± 1.47 kJ mol^-1^ with the energy minimum located at 1.72 nm. In contrast, SQ presented a ΔG value -22.78 ± 1.05 kJ mol^-1^ with the energy minimum positioned at 1.64 nm. Furthermore, an entry free energy barrier (ΔG^‡^) of approximately 4.7 kJ mol^-1^ at 2.64 nm for TRO and 1.5 kJ mol^-1^ at 2.58 nm for SQ was observed. SQ was therefore observed to penetrate deeper into the plasma membrane, indicating a higher affinity (more negative ΔG) compared to TRO. This ability was intuitively related to its molecular structure, which includes one less amine group than TRO (Figure 1). The lower charge on the SQ tail likely resulted in a reduced preference to interact with the aqueous environment. These variations could potentially be linked to the distinct polar:apolar balance values of each molecule.

**Figure 5:**
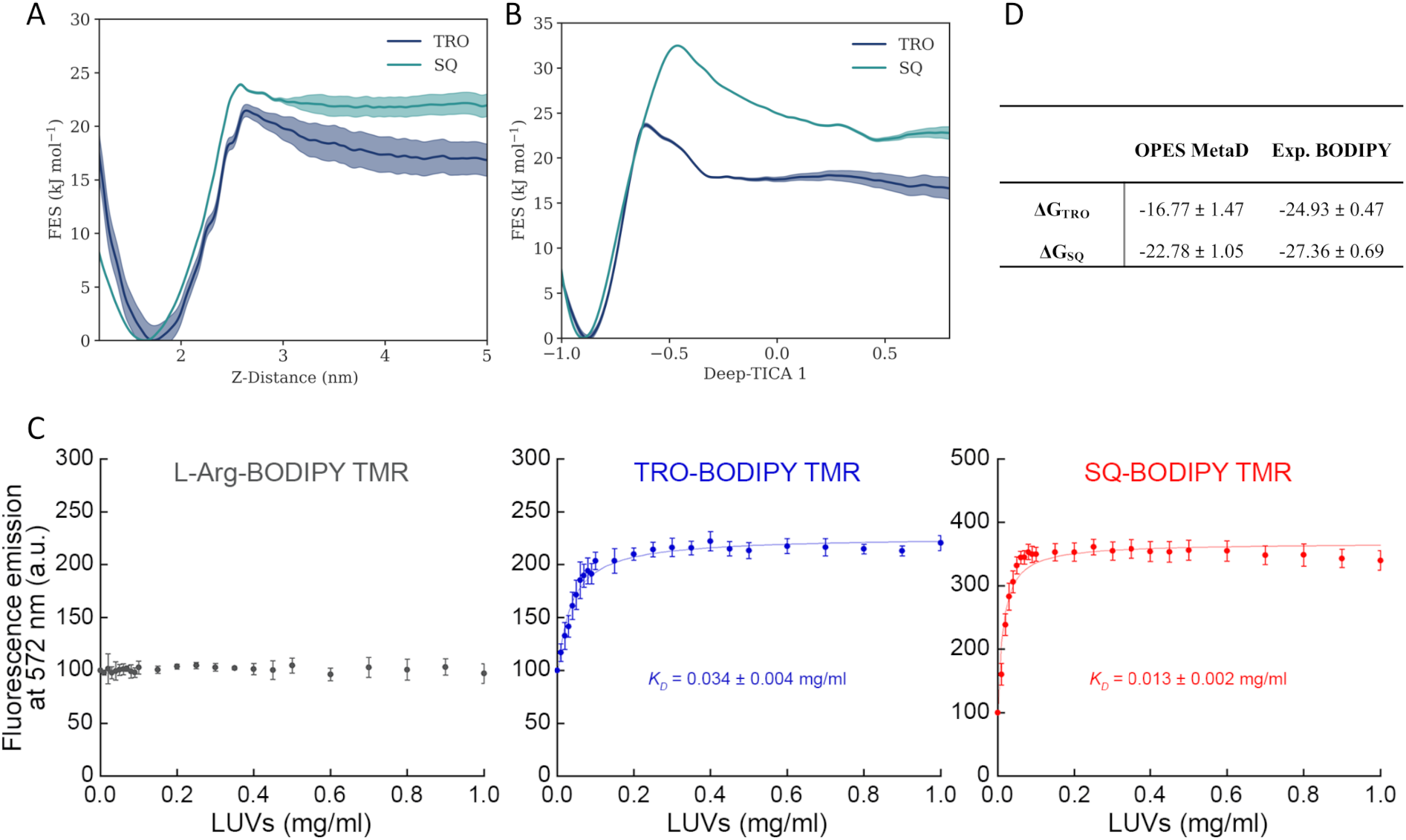
Aminosterols binding affinities. A) FES of TRO and SQ along the distance between the center of membrane and the aminosterols. B) FES of TRO and SQ along the Deep-TICA 1 CV. C) Binding of TRO and SQ to LUVs composed of 59% DOPC, 30% SM, 10% CHOL and 1% GM1. Binding plots reporting the fluorescence emission at 572 nm of 10 μM BODIPY TMR-labelled TRO (blue), SQ (red), and L-Arg (grey) versus LUV concentration. The lines through the data points represent the best fits to eq 4 (see experimental section). Each graph reports the obtained K_D_ value in units of mg/ml of total lipids. Experimental errors are SEM (n = 5). D) ΔG comparison between computational sampling and experimental binding data. All measures are expressed as kJ mol^-1^.

Membrane entry of aminosterols was observed to be a multidimensional phenomenon. Therefore, the slow entrance mechanism of TRO and SQ was better represented by the projection in the 2D plane of the Deep-TICA 1 component and the distance along the Z-axis, due to the intrinsic definition of the TICA variable (Figure S10, Supporting Informations). In this case, the Deep-TICA 1 CV allowed for a more accurate estimation of the FES (Figure 5B). The ΔG values were similar to those obtained with the previous analysis with the Z-distance as CV, obtaining -16.71 ± 1.1 kJ mol^-1^ for TRO and -22.75 ± 0.6 kJ mol^-1^ for SQ. This was consistent with the fact that the free-energy difference between the A and B states depends exclusively on the free-energy values at the two states, independently of the path that connects one state to the other. However, the ΔG^‡^ was approximately 7 kJ mol^-1^ for TRO and 10 kJ mol^-1^ for SQ. The observed discrepancy in the ΔG‡ identified along the Z-distance CV versus the Deep-TICA 1 CV is reflective of the dynamic that each variable represents. The Z-distance CV does not offer a precise energetic assessment of the membrane entry pathway, as it overlooks integral aspects such as aminosterols orientation during membrane insertion and their interactions with various lipid components. In contrast, the Deep-TICA 1 CV, formulated as a nonlinear combination of many variables, effectively discriminates between aqueous and lipid environments while accurately characterizing the orientation and lipidic interactions of aminosterols.

Notably, these energy barriers were surmountable in unbiased MD simulations dynamics^17^. Furthermore, the free energy minima identified by OPES aligned with the minima observed in classical MD simulations, demonstrating the consistency of our computational approach with the underlying physical phenomena^17^. Consequently, Deep-TICA 1 CV facilitated enhanced sampling among the metastable states of aminosterols and more effectively represented the energy barriers encountered along the membrane uptake pathway.

Furthermore, the advanced computational framework employed here delineated the membrane insertion process while also demonstrating exceptional proficiency in accurately estimating binding affinity within a reasonable simulation time. In order to evaluate the computational predictions, we estimated binding thermodynamics properties by quantitative measuring the affinity of BODIPY TMR labeled TRO and SQ labelled with BODIPY TMR for LUVs composed of 59% DOPC, 30% SM, 10% CHOL and 1% GM1, which is exactly the same lipid composition as that used for MD simulations. We performed binding experiments incubating 10 μM of each labelled aminosterol for 15 min with increasing concentrations of unlabeled LUVs. Both TRO and SQ displayed a significant increase in fluorescence emission in the presence of LUVs (Figure 5C), as previously observed^63^ and fitting the data points to a standard binding curve (Eq. 1) allowed us to obtain the corresponding *K*_*D*_ values (Figure 5C). The *K*_*D*_ values obtained with this LUVs composition were comparable to those obtained in a previous work with a very similar lipid composition, except a lower concentration of CHOL^63^. SQ was confirmed to have a higher increase of fluorescence upon LUV binding, and a higher affinity for LUVs, as showed by the lower *K*_*D*_ value of 0.013 ± 0.002 mg/ml (corresponding to a ΔG value of -27.36 ± 0.69 kJ mol^-1^), compared to 0.034 ± 0.004 mg/ml for TRO (corresponding to a ΔG value of -24.93 ± 0.47 kJ mol^-1^). The absolute ΔG values for SQ and TRO were within a discrepancy of 8 kJ mol^-1^, consistent with the typically force field errors of about ∼ 4–9 kJ mol^-1^ estimated in molecular dynamics (Figure 5D)^72,73^.

The computational framework introduced here offers versatility, allowing integration with other advanced sampling techniques like bias exchange, parallel tempering, or replica exchange. This integration could potentially lead to faster convergence, reducing computational time significantly. Moreover, the framework is adaptable, enabling the customization of descriptors based on the specific system under study, particularly focusing on the binding pockets of membrane proteins. Due to its versatility, this computational framework stands out as a highly valuable tool for tackling a broad range of challenges across chemistry, physics, and materials science.

## Conclusions

In the field of drug design and development research, understanding how drug candidates partition within cellular membranes is crucial. While recent experimental advances have provided valuable data on diverse compound-membrane interactions, the precision required to elucidate mechanisms of drug-membrane absorption—specifically, the orientation and location within membranes—remains a challenge. Such insights are critical for grasping the intricate molecular mechanisms underlying drug action. Computational methods offer supplementary perspectives that can complement and enrich this understanding. In this context, our study leveraged the synergy of enhanced MD simulations and innovative dimensionality reduction methods realized through the Deep-TICA approach. This methodology generated an effective CV specific of the molecular system under investigation, enhancing our ability to describe the drug absorption within membranes.

We tested the computational protocol on TRO and SQ, two aminosterols known for their ability to partially insert into neuron membranes, making them resistant to the toxicity of oligomers involved in AD and PD^39,42^. The results highlighted the complex, multidimensional nature of aminosterol membrane insertion, with two main metastable states corresponding to aqueous and lipid environments. The negative free energy difference between these states suggests a greater affinity of SQ and TRO for the neuron-like membrane, relative to the aqueous phase, indicating their preference for the membrane environment, as opposed to aqueous milieu. Such affinity also appears to be larger for SQ than for TRO, which can be rationalized by the shorter and less positively charged polyamine moiety in SQ. The free energy change values obtained from OPES MetaD aligned with experimental data obtained by fluorescence emission techniques, lending credibility to the findings.

Overall, the study demonstrated the effectiveness of the advanced computational framework employed in accurately estimating binding affinities and in contributing to a deeper understanding of aminosterol-lipid membrane interactions. In conclusion, the computational framework employed in this research holds great promise not only for furthering our knowledge of aminosterol compounds but also for advancing the broader field of drug discovery in the fight against neurodegenerative disorders and other complex disorders.

## Acknowledgments

This work was supported by a grant from the Swiss National Supercomputing Centre (CSCS).

## Notes

### Competing Interest Statement

The authors have declared no competing interest.

