## Supporting Information for "Leveraging Machine Learning-Guided Molecular Simulations Coupled with Experimental Data to Decipher Membrane Binding Mechanisms of Aminosterols"

### Unveiling Aminosterols Membrane Binding Mechanisms through the Integration of Experimental Data and Machine Learning Guided Molecular Simulations

Authors: Stefano Muscat[1], Silvia Errico[2], Andrea Danani[1], Fabrizio Chiti[2], Gianvito Grasso[1,*]

[1]Dalle Molle Institute for Artificial Intelligence IDSIA USI-SUPSI, Via la Santa 1, 6962 Lugano-Viganello, Switzerland;

[2] Department of Experimental and Clinical Biomedical Sciences, Section of Biochemistry, University of Florence, 50134 Florence, Italy

#### S1 - Trodusquemine and squalamine bond terms optimization.

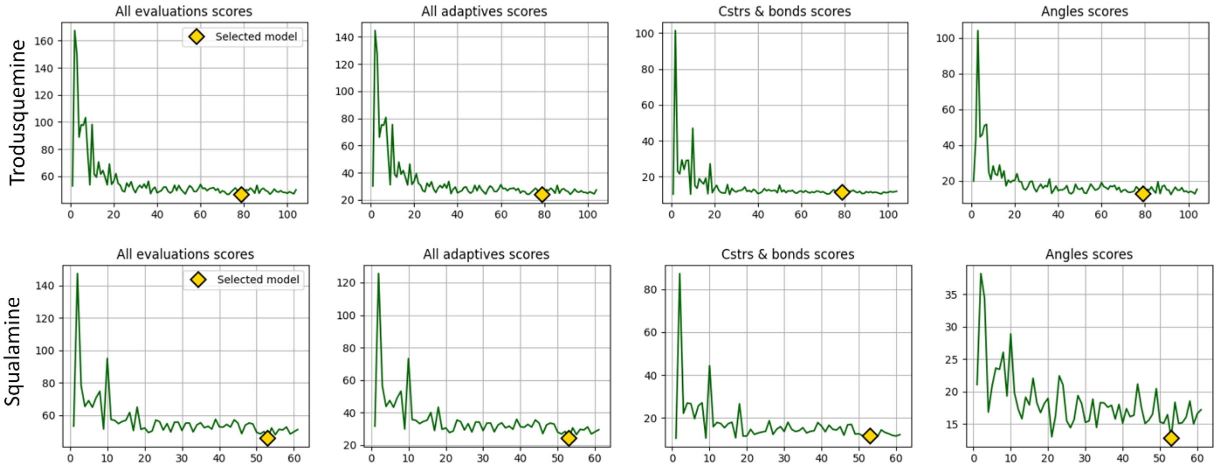


*Figure S1: Total optimization scores over iterations.*

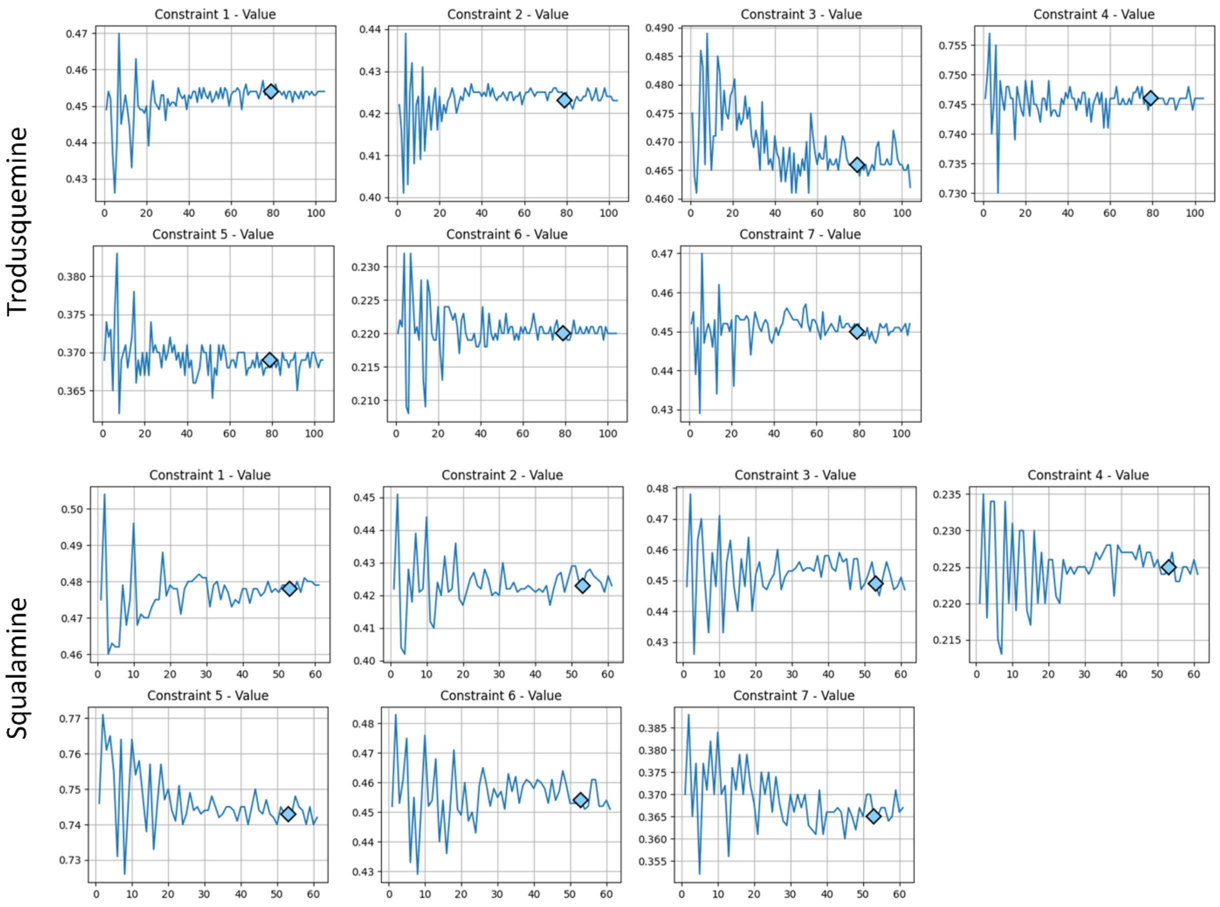


*Figure S2: Constraints optimization over iterations.*

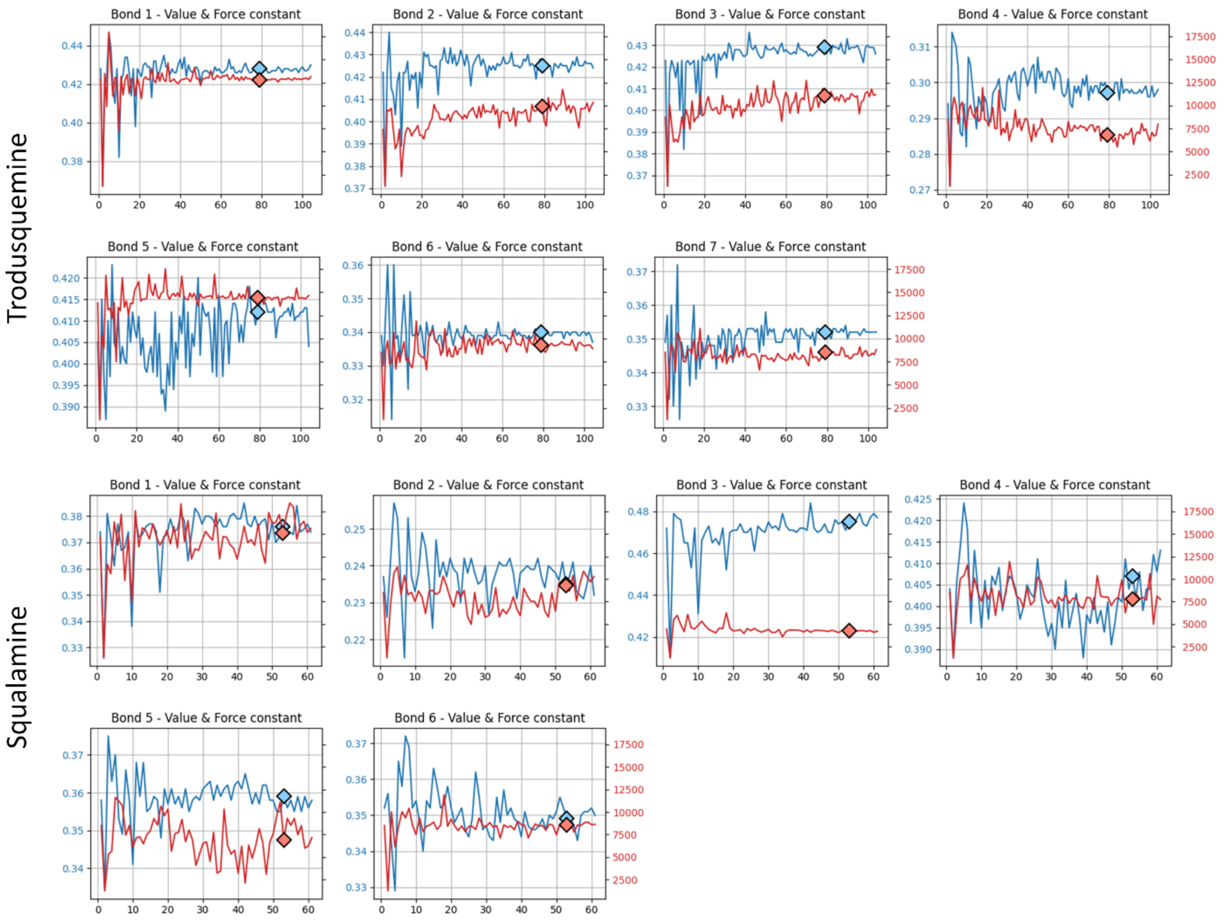


*Figure S3: Bonds optimization over iterations.*

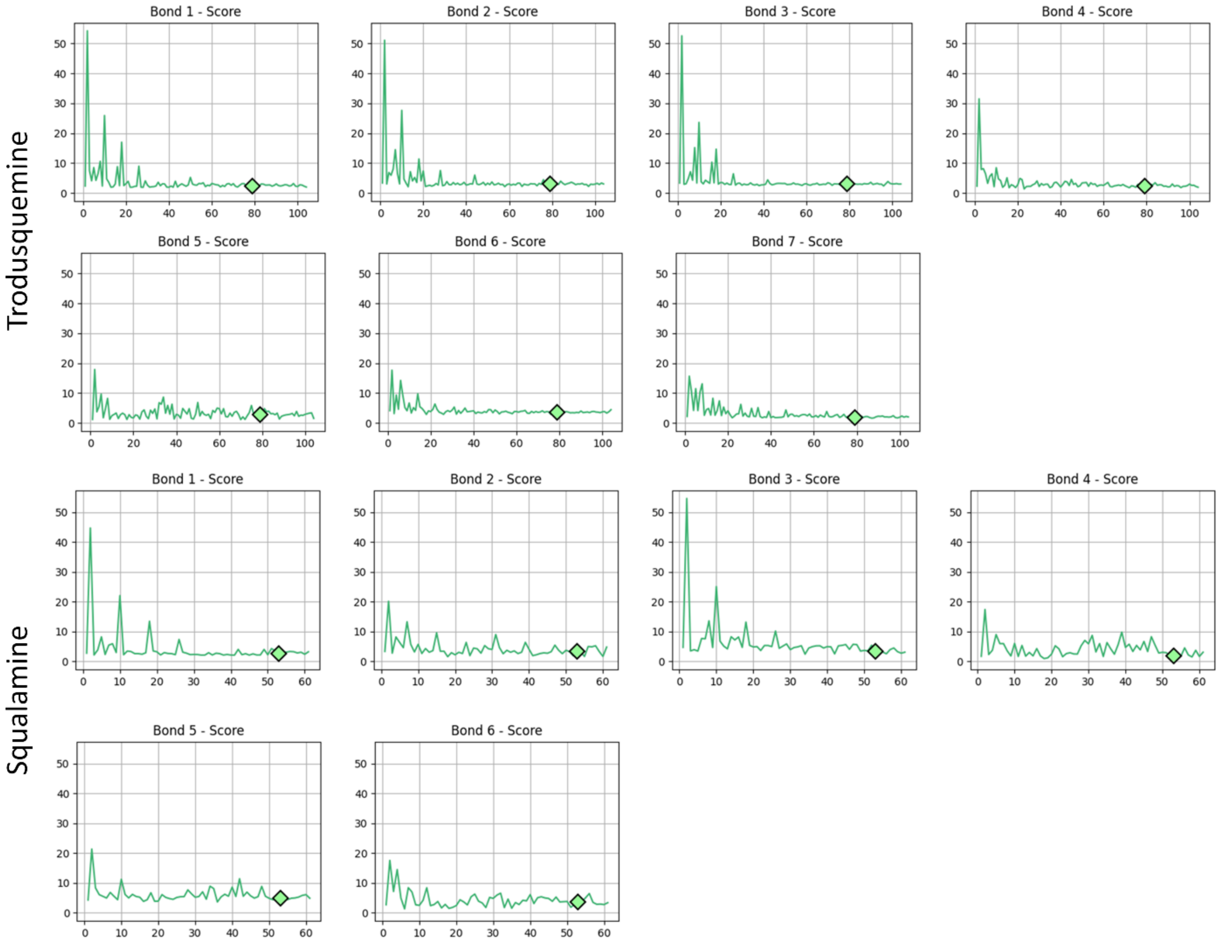


*Figure S4:Summary of the bond scores over iterations.*

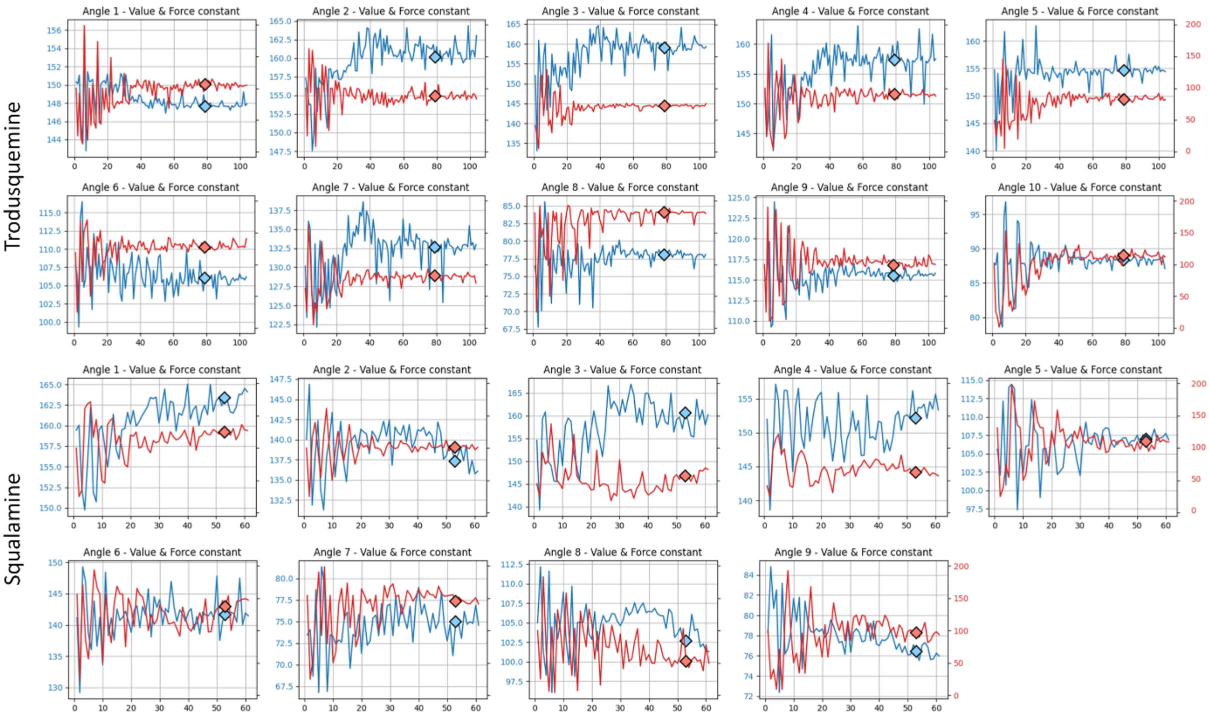


*Figure S5:Angles optimization over iterations.*

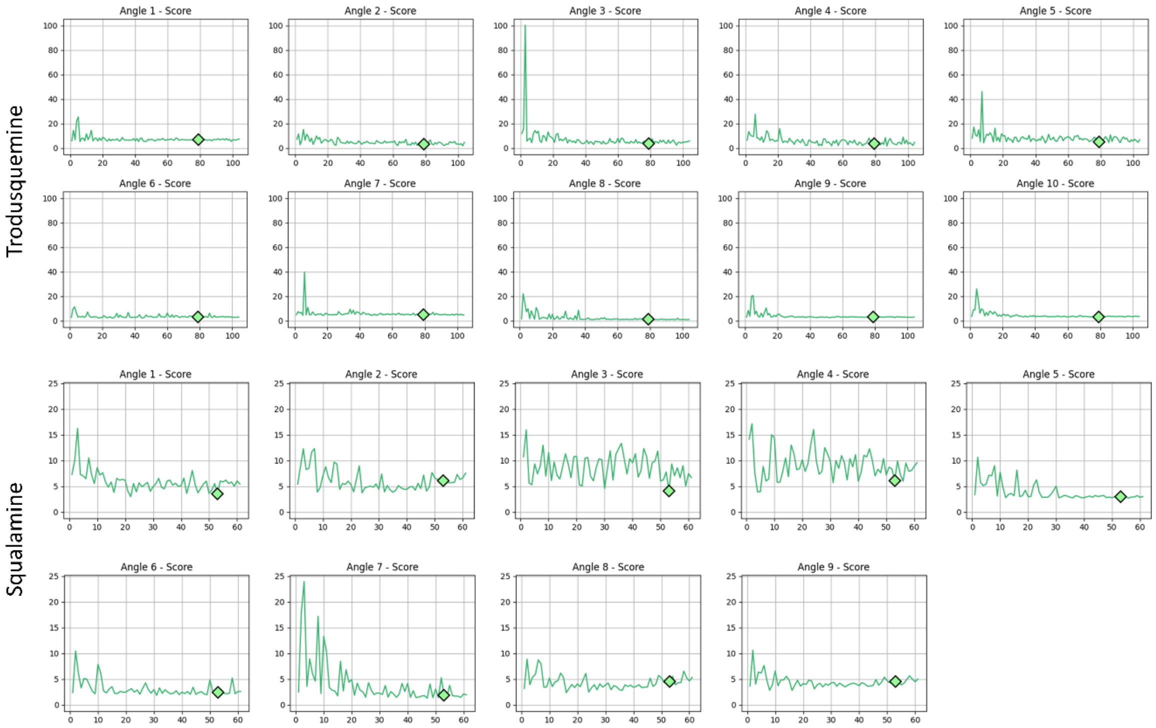


*Figure S6: Summary of angles over iterations.*

#### S2 – OPES multithermal convergence

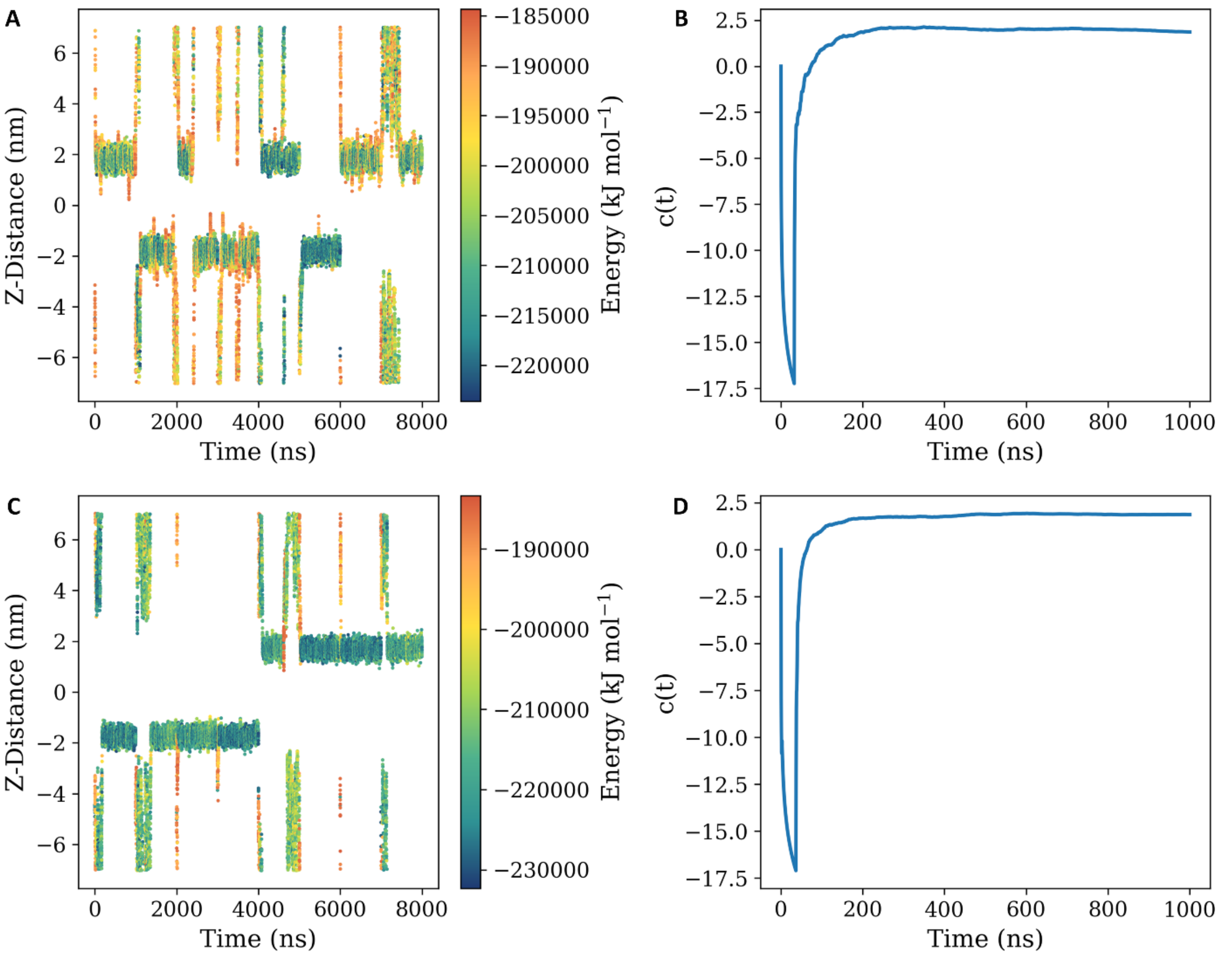


*Figure S7: Time evolution of the normal distance between the center of mass of aminosterol (trodusquemine in panel A and squalamine in panel C) and the center of the cell membrane, colored by the potential energy of the molecular system. Time evolution of the OPES multithermal bias shared between 8 replicas that asses the convergence of the simulation of trodusquemine (panel B) or squalamine (panel D) system.*

#### S2 – Deep-TICA training and validation

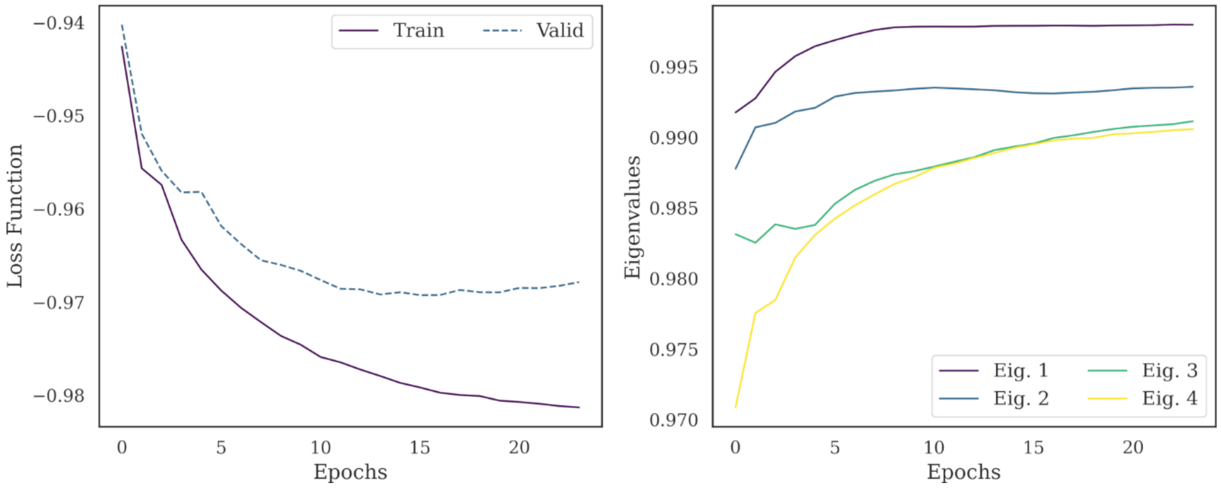


*Figure S8: Loss function for TICA training (left panel) and eigenvalue of first TICA vectors (right panel) vs epochs of trodusquemine system.*

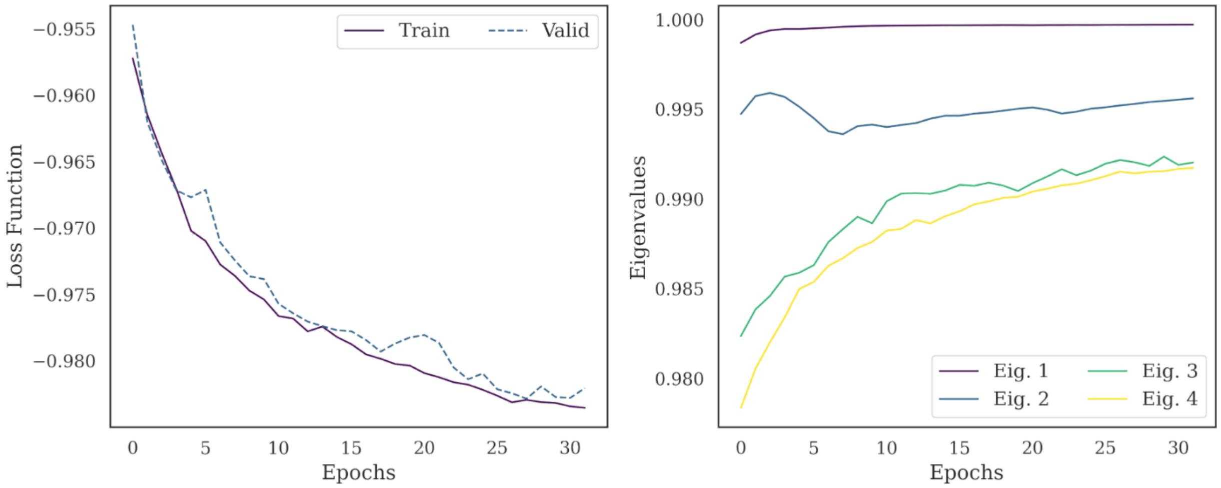


*Figure S9: Loss function for TICA training (left panel) and eigenvalue of first TICA vectors (right panel) vs epochs of squalamine system.*

#### S3 – FES 2D

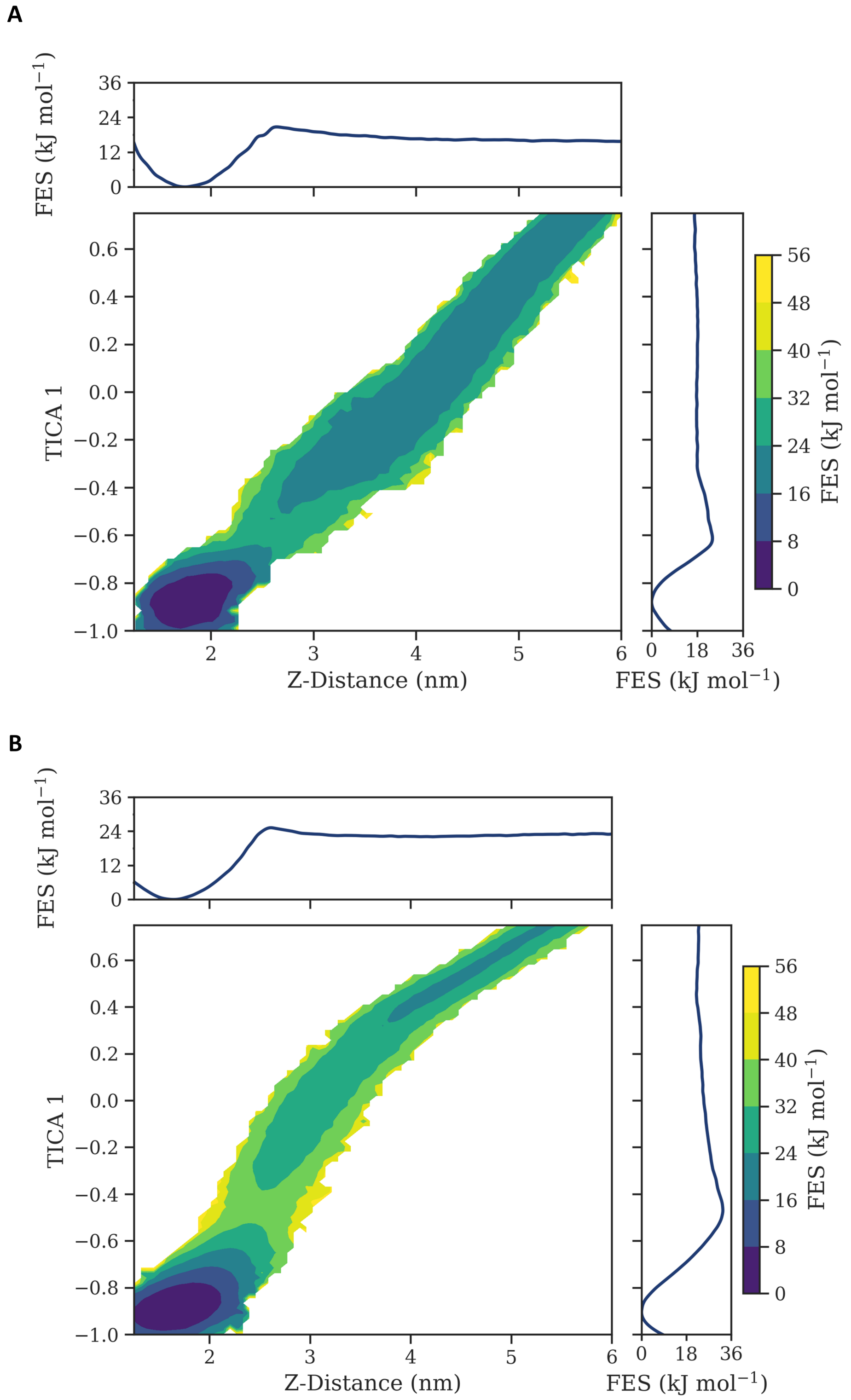


*Figure S10: The FES of TRO (panel A) and SQ (panel B) is illustrated as a function of the z-distance from the membrane center and the Deep-TICA 1 CV. The FES projections along the corresponding axis are displayed in the upper and right panels.*
